# Unequal contributions of species’ persistence and migration on plant communities’ response to climate warming throughout forests

**DOI:** 10.1101/217497

**Authors:** Romain Bertrand

## Abstract

Community reshuffling is lagging behind climate warming for many taxa, thereby generating a climatic debt. However, only few studies have attempted to assess the underlying factors that explain this debt, and none has gone further to explore this issue from a biogeographical perspective. Here I examine how effects of species’ migration and persistence on the current climatic debt vary spatially in forest herbaceous communities throughout the French territory. I show that Mediterranean communities are responding to climate warming through both high species’ migration and persistence effects, while alpine forest is the only ecosystem where species’ migration overtakes species’ persistence mechanisms. Such an approach seems promising in assessing the underlying mechanisms of the biodiversity response to climate change locally, and it can be applied for conservation issues to assess biodiversity sensitivity and optimize its management.

---

The concept of climatic debt or lag has been recently brought up to date in ecology (e.g. Menéndez et al. 2006) since its past development (Davis 1986). This debt assesses the time-delayed response of biological entities (e.g. species, communities or ecosystem) to climate change as a result of extinction debt and immigration credit (Jackson and Sax 2010). Studies have demonstrated that species composition in forest plant (Bertrand et al. 2011), butterfly, bird (Devictor et al. 2012) and freshwater fish (Comte and Grenouillet 2015) communities lagged behind the current climate warming. Recently, one step further has been taken by analyzing the underlying factors that explain this lag (Bertrand et al. 2016, Gaüzère et al. 2017). It has notably been shown that factors involved in species’ persistence are absorbing more of climate warming in communities than those involved in species’ migration are able to mitigate through thermophilization of forest plant assemblages (Bertrand et al. 2016). Such rare studies assumed this determinism was stable over large and heterogeneous areas. However, it is likely to vary locally following the proportion of warm-adapted species present in the regional species pool as well as the magnitude of the climate change or the habitat connectivity and disturbance for instance (Bertrand et al. 2011, 2016).

Based on this conceptual framework, I examined the spatial nonstationarity of species’ persistence and migration effects on the current climatic debt observed in French forest plant communities using Geographically Weighted Regression (see Supplementary material Appendix 1 for a data and model description, and Appendix 2 for a discussion on the model validity and outputs). The factors involved in species’ persistence accounting for in this analysis are tolerance to climatic stresses, temporal climatic niche shift (resulting from evolutionary adaptation, acclimation, and/or phenotypic plasticity), species longevity, nutrient resources, and microclimate buffering, while those involved in species’ migration include climatic niche tracking, habitat connectivity, earliness of seed dispersal, and competition for water resources (contributing to select warm-adapted species in plant communities) (Essl et al. 2015, Bertrand et al. 2016).

I show that the effects of species’ migration and persistence highly vary throughout the French forest territory (Figs. 1A and B). The magnitude of both effects is greater in the southern than the northern part of France. The plant communities in Mediterranean forest are intensely responding to the current warming and display the highest effects of species’ persistence and migration (Figs. 1A, B and D). This result likely corroborates the observed low sensitivity of these plant communities facing climate change (e.g. Vennetier and Ripert 2009) while other global change drivers can alter their sustainability (such as habitat loss and disturbance through fire and human actions; Sala et al. 2000). Mediterranean plants have developed efficient adaptations to hot and dry conditions, which allow them to absorb part of the climate change and hence to persist in communities (e.g. Thuiller et al. 2005). By contrast, the high species’ migration effect depicts a facilitated climatic niche tracking due to both rugged topography and proximity to Alps mountains that help species to find short-distance climatic escape leading to communities’ thermophilization and high species turnover (e.g. Thuiller et al. 2005).

**Figure 1:**
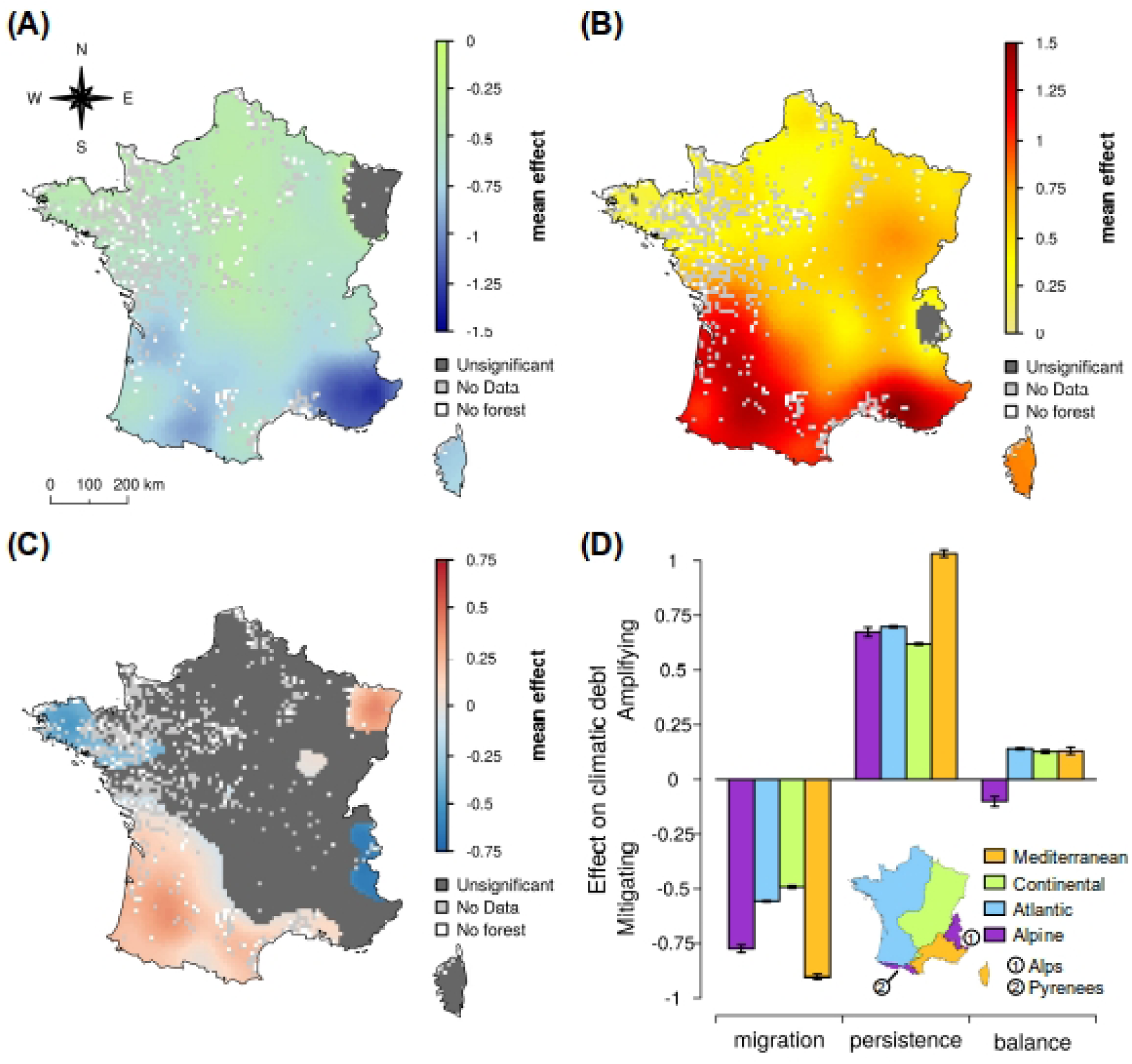
Spatial variation of the mean effects of migration (A) and persistence (B) mechanisms on the current climatic debt observed in plant communities throughout the French forests, and balance between both effects (C). (D) Synthesis of species’ migration and persistence effects in each biogeographic region encountered in France. Mean effects (and confidence intervals in panel D) are computed from 1000 bootstrapped models. More negative and positive values indicate higher mitigating and amplifying effects on the climatic debt, respectively.

Furthermore, I show that alpine plant communities are the only communities of the four biogeographic regions observed in France with a species’ migration effect taking over the species’ persistence effect (Figs. 1C and D). Climatic niche is more easily tracked by species in highland than in lowland areas (median isotherm shift since 1960 = 35.6 and 1.1 km in lowland and highland areas, respectively, according to Bertrand et al. 2011), hence reducing the climatic debt (60.7% of temperature increase recovered in highland forests, according to Bertrand et al. 2011). However, such a pattern varies inside the alpine biogeographic region itself with a higher effect of species’ persistence in Pyrenean forests (Fig. 1C). This contrasted determinism between Alps and Pyrenees mountains is likely explained by the proximity of the Mediterranean region to Alps mountains, which facilitates colonization of highlands by warm-adapted (sub-) Mediterranean plants, leading to high community thermophilization (Bertrand et al. 2016).

In conclusion, I emphasize high spatial disparity in species’ persistence and migration effects driving the forest plant community response to climate change. Reproducing such an analysis on other taxa should allow to get new insights on the nature, magnitude, and efficiency of mechanisms that biodiversity is implementing in the face of climate change. Predictive modeling will benefit these advances by preferentially integrating the most important drivers (Svenning and Sandel 2013). This approach is also promising for biodiversity conservation because it allows the assessment of how biodiversity is responding to environmental changes. Here Atlantic and Continental forest plant communities are likely more sensitive to the current warming due to both large climatic debt (Bertrand et al. 2016) and lower species’ migration and persistence effects than Mediterranean and alpine communities (Fig. 1). Such kind of information will be useful in optimizing biodiversity management (e.g. through selecting best actions considering the mechanisms that drive the current biodiversity to climate change) and in evaluating the efficiency of conservation actions (by assessing how they affect the biodiversity response to climate change).

## Acknowledgements

I thank S. Blanchet and J. Cote for useful comments; M. Loreau and J. Lenoir for discussions on this topic; the French National Institute of Geographic and Forest Information (IGN FI) to provide me with the NFI database; all those have contributed to the EcoPlant, Sophy and NFI databases, and especially J.-C. Gégout, I. Seynave, H. Brisse, P. de Ruffray, and F. Morneau. EcoPlant database was funded by the French Institute of Agricultural, Forest and Environmental Engineering (ENGREF, AgroParisTech), the National Forest Department (ONF), and the French Agency for Environment and Energy Management (ADEME). This study was supported by the TULIP Laboratory of Excellence (ANR-10-LABX-41).

## Appendix 1

### MATERIAL AND METHODS

#### Input data

All the data used in the present work have been already published. A detailed presentation and discussion of these data are provided in Bertrand et al. 2011 and 2016.

##### Floristic data

Plant community observations (from which the climatic debt is assessed) are investigated from 67289 georeferenced and dated floristic surveys recorded during the period 1987–2008 and stored in the French National Forest Inventory (NFI; Robert et al. 2010), Sophy (Brisse et al. 1995) and EcoPlant (Gégout et al. 2005) databases.

##### Environmental and biological determinants of the climatic debt

The effects of a set of 23 factors have been tested as in Bertrand et al. (2016). These abiotic, biotic and anthropogenic factors can be ordered considering their impact on the plant community response in the face of climate change (Dickinson et al. 2014, Essl et al. 2015, Bertrand et al. 2016).

First, eight factors have been selected to depict species’ persistence facing climate change in plant communities (Bertrand et al. 2016). Species’ intrinsic ability to tolerate climate change has been inferred as the temperature and precipitation niche breadths (i.e. the difference between the minimum and maximum annual mean temperature and annual precipitation where the species occurred). Both measures were then averaged across all species co-occurring within a given plant assemblage to get two community mean values characterizing the focal plant assemblage’s tolerance to thermal (*TO*_*T*_) and water (*TO*_*W*_) stresses. Species may also persist through evolutionary adaptation, acclimation, and/or phenotypic plasticity by shifting their ecological requirements (that is, niche shifts), to survive to the new climate conditions. Such biological factors were summarized in an index measuring the thermal niche shifts over the time considering any temporal changes in sampling effort (Bertrand et al. 2016), and then averaged among all species co-occurring in observed plant communities (*DC*). Species longevity (which increases the species resistance/tolerance to climate change and promotes community stability; Davis 1986) were collected from the LEDA database (Kleyer et al. 2008) and averaged among all plants co-occurring in observed communities (*LG*). Soil conditions defined here by soil acidity (*pH*) and N-nutrient availability (*N*) (bio-indicated from plant assemblages in absence of direct soil measures in every floristic surveys; Riofrío-Dillon et al. 2012, Riofrío-Dillon 2013) have been used to depict their alleviating effects of climatic constraints on plants (Bertrand et al. 2016). Microclimate buffering, due to local topography change (*THET*; e.g. Scherrer and Körner 2011) and canopy cover (*TBUF*; e.g. *Lenoir et al. 2017*) provide short-distance climate escapes and cooler conditions at the forest floor, respectively, contributes to local species’ persistence facing climate change. *THET* was inferred as temperature heterogeneity within each 1 km² unit (that is, the spatial resolution of all climatic grids I used due to the 500 m uncertainty in the geographic location of the floristic surveys) by using a finer-resolution temperature grid (2,500 m²; Bertrand et al. 2016). *TBUF* was computed from the *microclim* model (Kearney et al. 2014) as the difference between the temperatures perceived on the forest floor considering and not considering the observed canopy cover. All these factors allow plant communities to absorb part of the climate warming leading to an increase in the climatic debt (Bertrand et al. 2016).

Second, five factors have been used to depict species’ migration facing climate change (Bertrand et al. 2016). Climatic niche tracking (which underlies poleward and upward range shifts in response to climate warming) was inferred for each species as the similarity observed between the current thermal niche and the one expected if climatic niche tracking was maximal in data (Bertrand et al. 2016). Values of species co-occurring in plant communities were then averaged (*NC*). Both proximity to past species’ habitat (*HP*) and temporal changes in species’ habitat aggregation (*dHA*) have been computed for each species to define habitat connectivity. Both variables have been computed from temporal changes in thermal (defined as the projection across the geographical space of the realized thermal niche computed from data) and forest (derived from spatio-temporal Corine Land Cover layers) habitats, and then averaged considering the plant composition in communities. Competition for water resources, which contributes to select warm-adapted species in plant communities and hence favors their migration (Bertrand et al. 2016), was included in the analysis. It was computed for each species as the mean value of the hydric niche differentiation (i.e. *1-D* with *D* is the Schoener’s *D* index assessing niche overlap; Schoener 1968) among plants co-occurring in observed communities (*C*_*W*_). A high niche differentiation among co-occurring species depicts a high resource partitioning likely driven by a strong resource competition in a plant community (Bertrand et al. 2016). Species longevity is inversely correlated to species’ migration under the assumption that short-lived species have an earlier access to reproduction and hence to migration than the long-lived one. As a consequence, such a effect was also accounting for to depict species’ migration in the analysis. All these factors allow plant communities to mitigate part of the climate warming by promoting thermophilization of their plant assemblage, and as a consequence lead to reduce the climatic debt (Bertrand et al. 2016).

Third, 11 factors have been used to describe several dimensions of the environment from baseline environmental conditions and environmental changes. Climate conditions were extracted from spatio-temporal layers (Bertrand et al. 2011, 2016, Bertrand 2012; 1 km^2^ of spatial resolution). Baseline climate conditions were computed as the annual mean temperature (*T*) and the annual precipitation (*P*) over the 1965–1986 baseline period. Plant community exposure to climate change were computed as changes in annual mean temperatures (*TC*) and annual precipitations (*PC*) between the year of the floristic observations and the 1965–1986 baseline period. The direct impact of light availability (*L*) on the composition of understory plant communities (through disturbing resident communities as light increases sharply; e.g. Wagner et al. 2011) was inferred by computing the average value of the L-Ellenberg index of each species co-occurring in communities (Ellenberg et al. 1992). Competition for soil nutrients (*C*_*N*_), competing with climate change effects (Bertrand et al. 2016), was inferred as the mean value of N-nutrient niche differentiation (using the same method as the one used to compute *C*_*W*_; see above) among plants co-occurring in observed communities. Anthropogenic and natural disturbances were inferred by five more variables: the presence/absence of recent silvicultural practices (*SILVP*), the presence/absence of human-mediated and natural disturbances (*DISTURB*) and the presence/absence of exotic trees (*EXOT*) extracted from the NFI database, as well as the proximity to road (*RP*; ranging from forest path to highway, computed from the GEOFABRIK spatial layers) and the human population density (*HPD*; extracted from the Insee database). The effects of all these variables are not investigated in the present study, but are used to fix the effects of species exposure to global changes in the model. Such factors are known to interact in or impact directly the biodiversity response to climate change without reflecting any species’ persistence or migration mechanisms. They depict environmental pressures that inflate the climatic debt through community reshuffling towards another environmental equilibrium (compared to the climate one) or mortality (Bertrand et al. 2016).

Although several variables were computed from bioindication methods (*L*, *pH*, *N* and the floristically reconstructed temperature used to infer the climatic debt), they were largely uncorrelated (R²<0.1), demonstrating no circularity issues between these indices.

#### Inference of the climatic debt

The climatic debt was not computed in the present study but existing values were analyzed from Bertrand et al. (2016). The climatic debt (*dT*) was computed as the difference between the annual mean temperature at a given location and year, and the annual mean temperature of the same location and year but inferred from plant assemblages (*FrT* as floristically reconstructed temperature) (Bertrand et al. 2011, 2016). A positive difference means that the observed reshuffling in a plant community is lagging behind climate warming (Bertrand et al. 2011), and thus depicts a climatic debt for that species assemblage (Devictor et al. 2012). *FrT* values were modeled in a previous studies (Bertrand et al. 2011) using a transfer function that combines a weighted averaging *PLS* regression (accounting for linear effect; ter Braak and van Dame 1989) and a Breiman’s random forest model (accounting for nonlinear and interaction effects among species occurrences in residuals of the weighted averaging *PLS* model; Breiman 2001) to infer temperatures from plant community composition (presence/absence of a set of 760 herbaceous species). This approach was validated on an independent data set of 5136 floristic surveys (R²=0.83, RMSD=0.97 °C; Bertrand et al. 2011). Floristic surveys used to adjust and valid the model came from the NFI, Sophy and EcoPlant databases.

#### Analytical approach

I have employed the same approach that it has been conducted in Bertrand et al. (2016), but I used a different and appropriate statistical method to test for spatial nonstationarity in the climatic debt determinism. The analysis was conducted by fitting Geographically Weighted Regression (*GWR*; Fotheringham et al. 2002). The hypothesis behind this particular regression was that the fitted coefficients of a global model (i.e. fitted to all the data as in Bertrand et al. 2016) may not represent local variations in the data (Bivand et al. 2008). *GWR* explored spatial nonstationarity by moving a weighted window over the spatial distribution of data, estimating one set of coefficient values at every observation from the adjustment of a local linear model (Bivand et al. 2008).The weighted window was determined by cross-validation searching for the proportion of observations to include in the weighted scheme that minimized the root mean square error of predictions (Bivand et al. 2008), and fixed to the 100 closest observations of each floristic survey present in the sample. This set of observations is weighted using a Gaussian spatial weighting function in order to give more weight to the data closed to the focal floristic observation in the local model adjustment.

The GWR model that was fitted is:

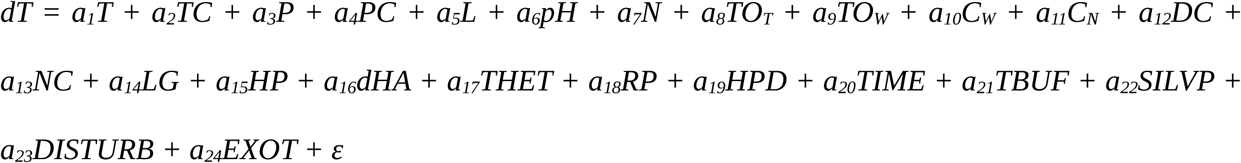

where *a*_1,…,24_ are the local estimated coefficients which I used to quantify the magnitude of the effect of each variable (see the text above for a definition and description of each variable as well as Bertrand et al. 2016 for more details), and *ε* denotes the residuals. All variables were centered and reduced to allow parameter comparison.

The *GWR* model was adjusted using a sub-sampling approach (*n* = 1000) in order to compute robust and accurate parameter estimation and uncertainty. Subsamples were composed of a set of 4830 floristic observations selected randomly (with no replacement) among 45806 floristic surveys (all coming from the NFI database since 1993) and distant 10 km from each other. Such a distance criterion aims to limit the effect of spatial autocorrelation in the model (Bertrand et al. 2016). Floristic observations in each of the 1000 subsamples were weighted in the *GWR* model by the inverse of the total number of observations per year in order to correct for an unbalanced temporal distribution of samples. The local coefficients of factors involved in species’ persistence and migration mechanisms were summed in order to assess their respective effects on the climatic debt in each of the 1000 subsamples. I tested the significance of these cumulative effects locally by comparing the distribution of their 1000 local coefficient values to 0. I considered that significant negative and positive coefficient values have at least 95% of the 1000 coefficients values less and more than 0 (i.e. a bootstrap test with a threshold α = 0.05), respectively. Finally, I mapped the effects of species’ persistence and migration, and compared their magnitude between biogeographical regions.

Analyses were conducted using the *R* freeware (R Core Team 2017) and the *spgwr* (Bivand and Yu 2015) R-package to fit the *GWR* model.

## Appendix 2

### Analysis and discussion of the *GWR* results

#### Model quality

*GWR* explains between 34.7 and 77.9% of the total variation of the climatic debt (i.e. adjusted R² value expressed in %) throughout the French forests (Fig. A1). At the French scale, the mean adjusted R² value (0.607; Fig. A1B) is higher than the one reported previously using *PLS* regression which is a more global statistical approach (R^2^ = 0.413 from Bertrand et al. 2016). It is noteworthy that part of this difference in the explanation of the climatic debt between the two approaches is due to different statistical objectives. In fact, the *PLS* regression used by Bertrand et al. 2016 has a lower R² value than a simple linear model (for instance) due to the component selection which allows to fit the model on a limited but clean part of the climatic debt variation that the set of explanatory variables can explain considering potential noises or biases presents in the data (such as multicollinearity for instance). However, these results demonstrate the high quality of the *GWR* adjustment and the high confidence that we can have in the species’ persistence and migration effects reported in the present study. *GWR* improves the explanation of the determinism of the climatic debt and shows the importance to account for spatial variability when the process is modeled from a database covering a large area (notably due to changes in ecological context and/or species pool that occur at local or regional scales).

**Figure A1:**
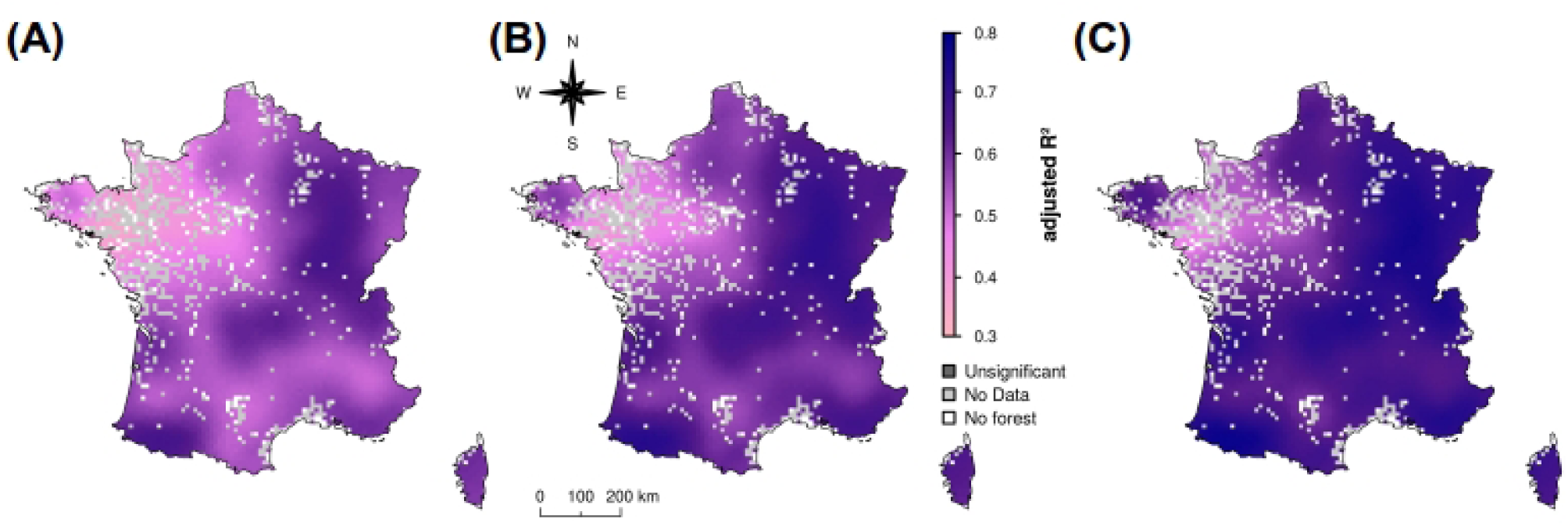
Map of adjusted R^2^ inferred from *GWR* (10 × 10 km^2^ spatial resolution). Lower (A) and upper (C) boundaries of the 95% confidence interval, and mean estimate (B) are mapped and computed from 1000 bootstrapped models.

Areas with the lowest R^2^ values (as in northwestern France) are areas where the set of explanatory factors fails to explain the climatic debt. It means that other ecological, biological or anthropogenic factors drive the climatic debt in these areas or that proxies that I computed and used as explanatory variables failed to mirror ecological, biological or anthropogenic drivers (see Bertrand et al. 2016 for a discussion on the limitation of variables aiming to depict forest disturbances and management for instance). To get a more accurate understanding of the determinism of the climatic debt in these cases, it will need to conduct local or regional studies with relevant measures of the environment and ecological context at this scale.

#### Variable contribution

Despite that it is not the aim of the present study, I provide additional results about the contribution of each factor involved in species’ persistence and migration mechanisms that I have tested in the *GWR* model. The contribution has been computed as the coefficient values of each variable divided by the sum of coefficient values of the set of factors involved in species’ migration or persistence (mapped in Figs. 1A and B), and expressed in percent. It assesses the weight of each variable in the definition of the species’ persistence and migration effects on the climatic debt. I show that the contributions highly vary among the factors and throughout forests (Figs. A2 and A3).

**Figure A2:**
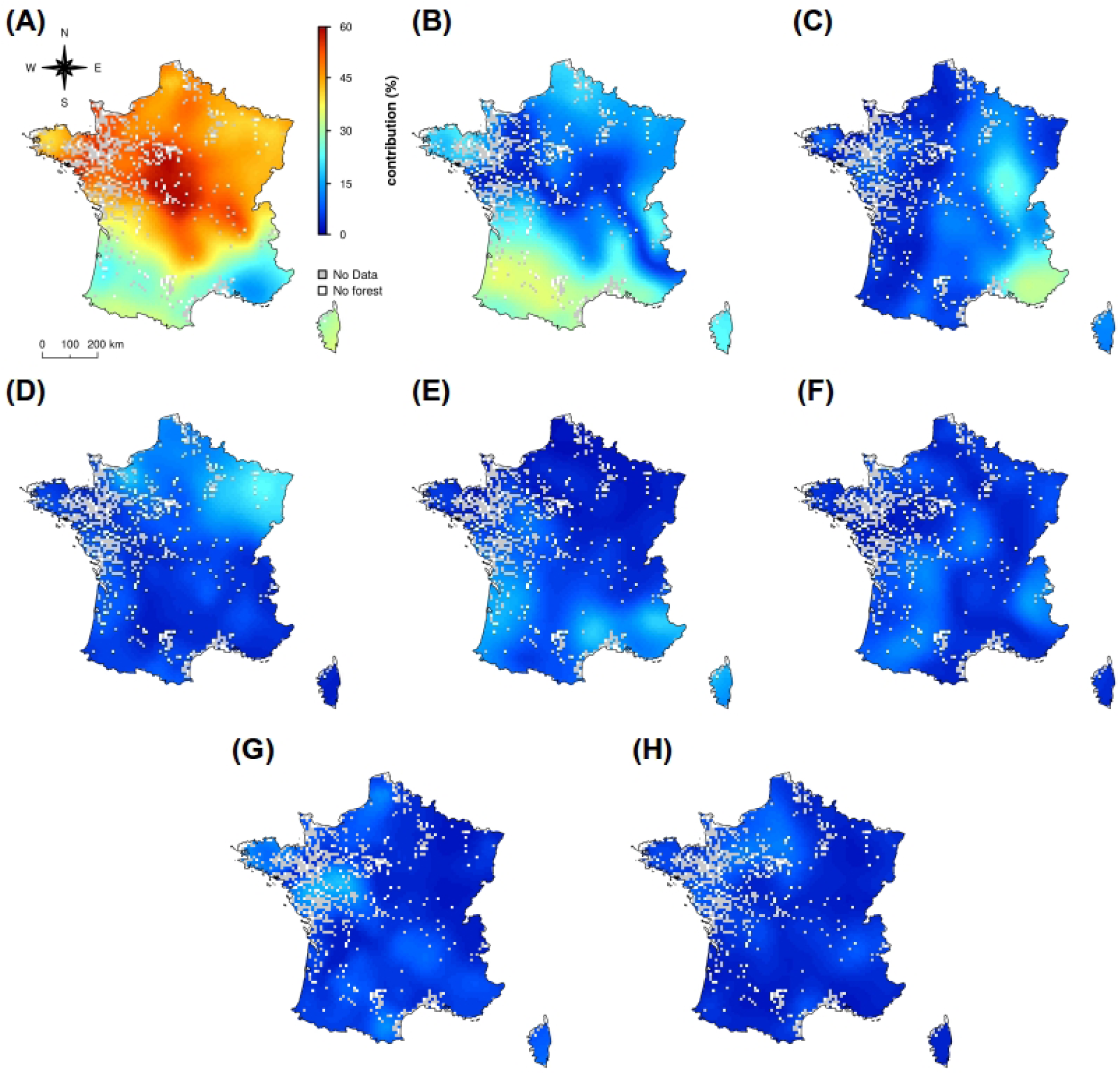
Map of the contribution of each ecological and biological factors involved in species’ persistence (10 × 10 km^2^ spatial resolution): species’ tolerances to hydric (A) and thermal stresses (B), climatic niche shift (C), pH (D), species longevity (E), N-nutrient availability (F), local temperature heterogeneity (G) and climate canopy buffering (H). Mean contribution estimates are mapped and computed from 1000 bootstrapped *GWR* models. The contribution of each variable is computed as the coefficient value of each of them divided by the sum of coefficient values of the set of factors involved in species’ persistence (i.e. the effect of species’ persistence in the determinism of the climatic debt reported in Fig. 1B), and expressed in percent.

**Figure A3:**
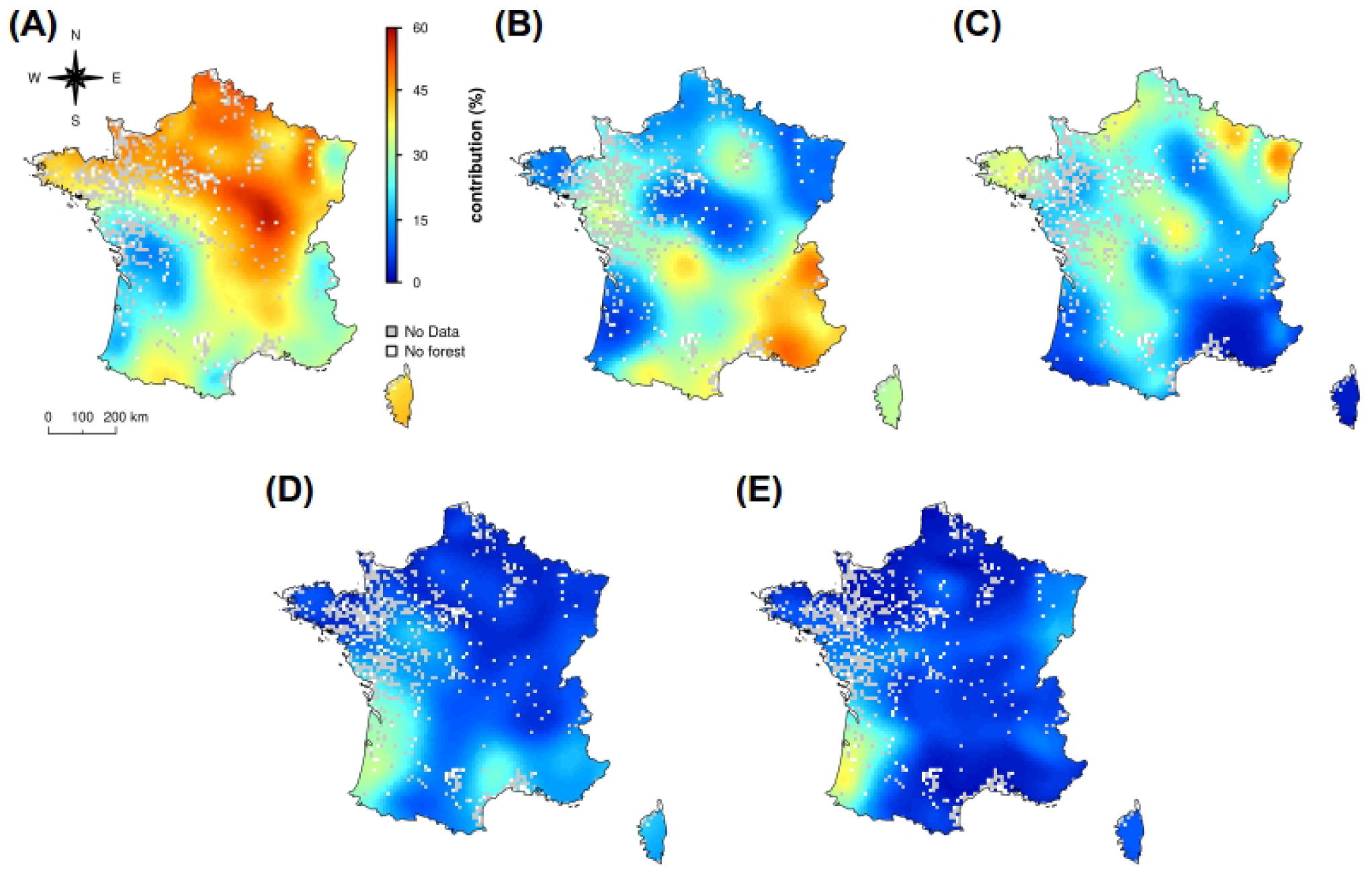
Map of the contribution of each ecological and biological factors involved in species’ migration (10 × 10 km^2^ spatial resolution): competition for water resources (A), climatic niche conservatism (B), temporal changes in species’ habitat aggregation (C), seed dispersal earliness (D) and proximity from past species’ habitat (E). Mean contribution estimates are mapped and computed from 1000 bootstrapped *GWR* models. The contribution of each variable is computed as the coefficient value of each of them divided by the sum of coefficient values of the set of factors involved in species’ migration (i.e. the effect of species’ migration in the determinism of the climatic debt reported in Fig. 1A), and expressed in percent.

Species’ tolerance to hydric stresses highly contributes to determine the species’ persistence effect on the climatic debt in the northern part of France (reaching 60% in some areas), while species’ tolerance to thermal stresses and climatic niche shifts (depicting adaptation, acclimation, and/or phenotypic plasticity effects) highly drive this determinism in the southwestern and southeastern part of France, respectively (Fig. A2). Such a spatial pattern is likely to demonstrate local or regional variations in hydric and thermal pressures on the forest flora. It also demonstrates that rapid climatic niche shifts occurred since 1993 in Mediterranean forests. However, it is difficult to state whether some traits of the Mediterranean vegetation and/or the increasing climatic stress observed in this region explains the high contribution of climatic niche shifts. Other factors contribute less than 20% to the species’ persistence effect on the climatic debt.

Species’ competition to water resources highly contributes to determine the species’ migration effect on the climatic debt in the northern part of France (reaching 60% in some areas; Fig. A3). This result as well as the high contribution of species’ tolerance to water stress reported above highlight water resources and its spatio-temporal variation as an important dimension of the climate change effect on forest plants in this area. It does not mean that water is a more limiting resource than in Mediterranean region, but that the species pool is likely less adapted by the recent changes in water regime in northern France than in the South (where the exposure to drought has selected highly adapted species to limiting water resource; e.g. Thuiller et al. 2005). Climatic niche tracking (depicting species’ migration towards suitable climate conditions) mainly drives the species’ migration effect in Mediterranean and alpine biogeographical regions (except in the western part of the Pyrenees; Fig. A3). The rugged topography encountered in these areas (promoting short-distance climate escapes) as well as the proximity between the Alps and the Mediterranean area (favoring exchange of species adapted to a large range of thermal conditions) are favorable conditions for climatic niche tracking. Temporal changes in species’ habitat aggregation also mainly contribute to determine the species’ migration effect on the climatic debt over smaller areas. Earliness of seed dispersal and proximity to past species’ habitat have low contribution (less than 20%) except in the southwestern part of France.

These results confirm the previous order of the most important factors involved in species’ persistence and migration established by Bertrand et al. (2016). However, they allow to move beyond the previous global findings (Bertrand et al. 2016) by highlighting some areas of transition among the effects of all the drivers. These results could serve of starting point of future detailed analysis aiming to verify the different plant response pathways to climate change reported here, and to understand what are the causes of this spatial structure.

#### Multicollinearity issue

*GWR* is often considered highly sensitive to multicollinearity (e.g. Wheeler and Tiefelsdorf 2005) while some recent results showed it is robust to its effects (Fotheringham and Oshan 2016). Multicollinearity between explanatory variables inflates the variance of regression parameters which potentially leads to both unstable model fit (due to high variance) and error in identification of relevant predictors as well as in their relative importance assessment (e.g. Dormann et al. 2013). It is still a big issue in statistics especially when real data varying in space and time are studied. No statistical models or methods are able to fully tackle this issue, but it is possible to assess whether multicollinearity is a matter of concern.

I computed correlation between explanatory variables, condition number computed from the eigenvalues of the model matrix (*CN*; e.g. Belsley et al. 2004), variance inflation factor (*VIF*; e.g. Belsley et al. 2004) in order to assess whether multicollinearity can alter my results and conclusions. First, explanatory variables are weakly correlated between them (Bertrand et al. 2016). The upper correlation value reaches a R^2^ of 0.381 which is lower than the accepted threshold of 0.49 (that Dormann et al. 2013 have shown to be an appropriate indicator for when collinearity begins to severely distort model estimation). Second, *CN* values (which assess the multicollinearity effect on the model fit) vary between 7.1 and 56.7 throughout the French forest territory (Fig. A4). *CN* exceed the accepted threshold of 30 (which identifies a potential severe multicollinearity concern; Belsley et al. 2004) in only rare areas such as small parts of the center (close to Paris) and the southwestern of France (covering a total of 10800 km^2^; Fig. A4B). Third, *VIF* values (which assess the sensitivity of each explanatory variable to multicollinearity) vary between 1.1 and 15.4 among the set of explanatory variables involved in species’ persistence and migration mechanisms throughout French forests (Figs. A5 and A6). *VIF* values exceed the accepted threshold of 10 (which identifies a potential severe multicollinearity concern for a variable; Belsley et al. 2004) for local temperature heterogeneity in a restricted part of the northern France (Fig. A5). All other factors have relatively low *VIF* values. These results demonstrate that multicollinearity is low in the present data and as a consequence its impact on the *GWR* model is unlikely altering my estimation of the species’ persistence and migration effects on the climatic debt as well as the conclusions of the study.

**Figure A4:**
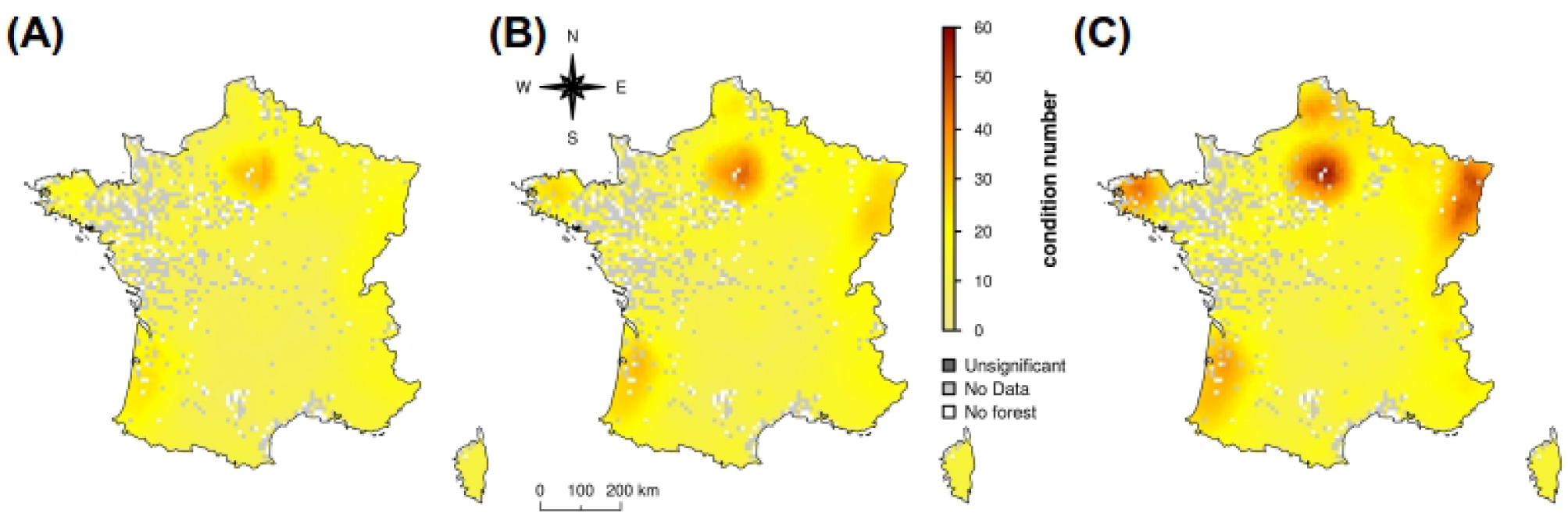
Map of the condition number (*CN*) inferred from *GWR* (10 × 10 km^2^ spatial resolution). Lower (A) and upper (C) boundaries of the 95% confidence interval, and mean estimate (B) are computed from 1000 bootstrapped models. A *CN* value more than 30 means that serious multicollinearity concerns can alter the model fit (Besley et al. 2004).

**Figure A5:**
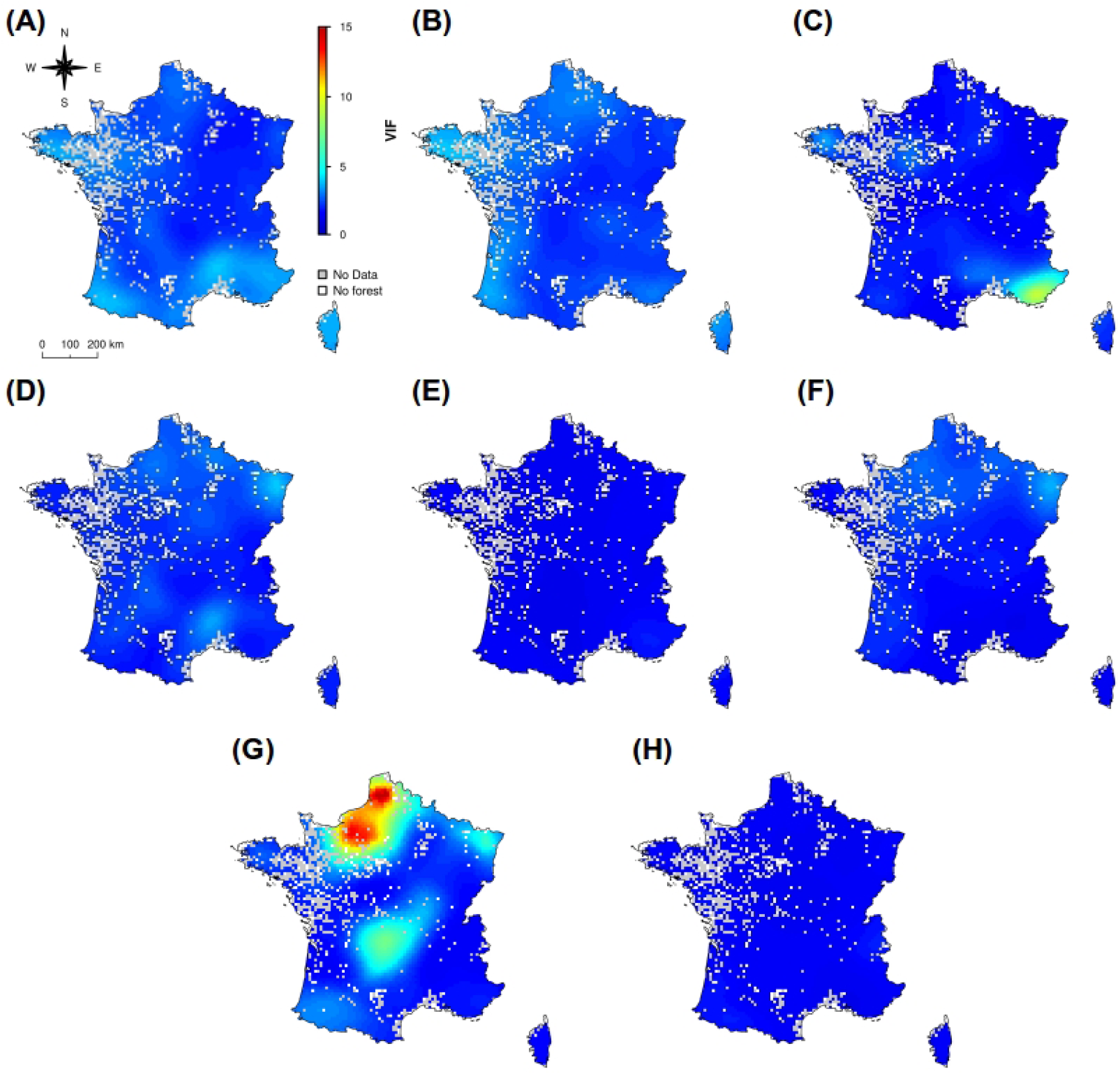
Map of the variance inflation factor (*VIF*) computed from *GWR* for variables involved in species’ persistence mechanisms (10 × 10 km^2^ spatial resolution): species’ tolerances to hydric (A) and thermal stresses (B), climatic niche shift (C), pH (D), species longevity (E), N-nutrient availability (F), local temperature heterogeneity (G) and climate canopy buffering (H). Mean *VIF* estimates are mapped and computed from 1000 bootstrapped models. A *VIF* value more than 10 means that the variable is potentially affected serious multicollinearity concerns (Belsley et al. 2004).

**Figure A6:**
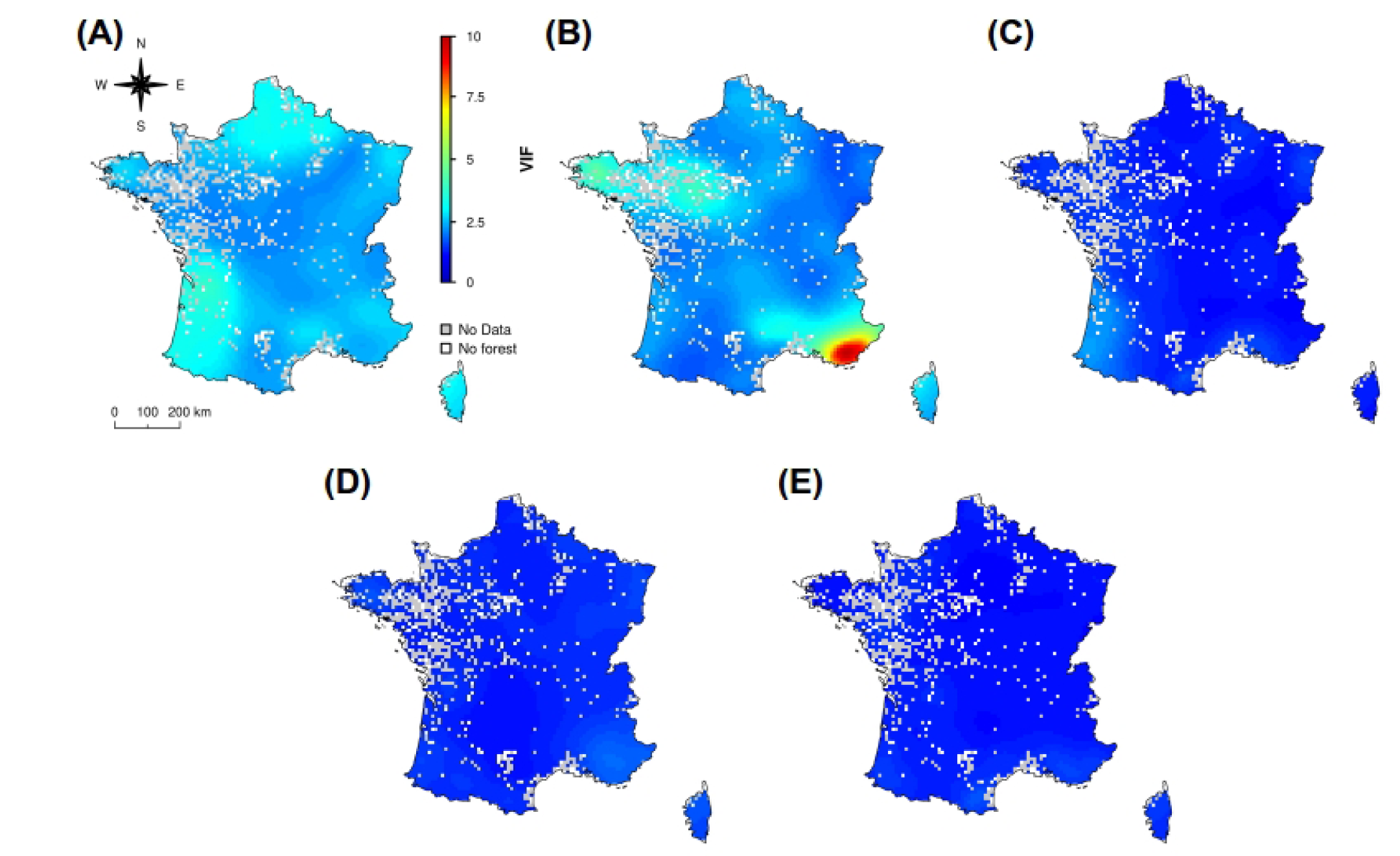
Map of the variance inflation factor (*VIF*) computed from *GWR* for factors involved in species’ migration mechanisms (10 × 10 km^2^ spatial resolution): competition for water resources (A), climatic niche conservatism (B), temporal changes in species’ habitat aggregation (C), seed dispersal earliness (D) and proximity from past species’ habitat (E). Mean *VIF* estimates are mapped and computed from 1000 bootstrapped model. A *VIF* value more than 10 means that the variable is potentially affected by serious multicollinearity concerns (Belsley et al. 2004).

## References

Bertrand, R. et al. 2011. Changes in plant community composition lag behind climate warming in lowland forests. – Nature 479: 517–520.

Bertrand, R. et al. 2016. Ecological constraints increase the climatic debt in forests. – Nat. Commun. 7: 12643.

Comte, L. and Grenouillet, G. 2015. Distribution shifts of freshwater fish under a variable climate: comparing climatic, bioclimatic and biotic velocities. – Divers. Distrib. 21: 1014–1026.

Davis, M. B. 1986. Climatic instability, time lags, and community disequilibrium. – In: Diamond, J. M. and Case, T. J. (eds), Community Ecology. Harper & Row, pp. 269–284.

Devictor, V. et al. 2012. Differences in the climatic debts of birds and butterflies at a continental scale. – Nat. Clim Change 2: 121–124.

Essl, F. et al. 2015. Delayed biodiversity change: no time to waste. – Trends Ecol. Evol. 30: 375–378.

Gaüzère, P. et al. 2017. Where do they go? The effects of topography and habitat diversity on reducing climatic debt in birds. – Glob. Change Biol. 23: 2218–2229.

Jackson, S. T. and Sax, D. F. 2010. Balancing biodiversity in a changing environment: extinction debt, immigration credit and species turnover. – Trends Ecol. Evol. 25: 153–160.

Menéndez, R. et al. 2006. Species richness changes lag behind climate change. – Proc. R. Soc. Lond. B Biol. Sci. 273: 1465–1470.

Sala, O. E. et al. 2000. Global biodiversity scenarios for the year 2100. – Science 287: 1770–1774.

Svenning, J.-C. and Sandel, B. 2013. Disequilibrium vegetation dynamics under future climate change. – Am. J. Bot. 100: 1266–1286.

Thuiller, W. et al. 2005. Climate change threats to plant diversity in Europe. – Proc. Natl. Acad. Sci. U. S. A. 102: 8245–8250.

Vennetier, M. and Ripert, C. 2009. Forest flora turnover with climate change in the Mediterranean region: A case study in Southeastern France. – For. Ecol. Manag. 258: S56–S63.

## References

Bertrand, R. 2012. Spatio-temporal response of the forest vegetation to climate warming – Assessment of the vegetation reshuffling and characterisation of the effect of ecological and geographical factors modulating this process at the species and community scales. – Ph.D. thesis, AgroParisTech.

Bivand, R. and Yu, D. 2015. spgwr: Geographically Weighted Regression. – R package ver. 0.6-28, <https://CRAN.R-project.org/package=spgwr>.

Bivand, R. et al. 2008. Applied Spatial Data Analysis with R. – Springer.

Breiman, L. 2001. Random Forests. – Mach. Learn. 45: 5–32.

Brisse, H. et al. 1995. European vegetation survey : la banque de données phytosociologiques “Sophy.” – Ann. Bot. 53: 191–222.

Devictor, V. et al. 2012. Differences in the climatic debts of birds and butterflies at a continental scale. – Nat. Clim. Change 2: 121–124.

Dickinson, M. G. et al. 2014. Separating sensitivity from exposure in assessing extinction risk from climate change. – Sci. Rep. 4: 1–6.

Ellenberg, H. et al. 1992. Zeigerwerte von Pflanzen in Mitteleuropa. – Scr. Geobot. 18: 1–258.

Fotheringham, A. S. et al. 2002. Geographically Weighted Regression: The Analysis of Spatially Varying Relationships. – Wiley.

Gégout, J.-C. et al. 2005. EcoPlant: A forest site database linking floristic data with soil and climate variables. – J. Veg. Sci. 16: 257–260.

Kearney, M. R. et al. 2014. microclim: Global estimates of hourly microclimate based on long-term monthly climate averages. – Sci. Data 1: 140006–.

Kleyer, M. et al. 2008. The LEDA Traitbase: a database of life-history traits of the Northwest European flora. – J. Ecol. 96: 1266–1274.

Lenoir, J. et al. 2017. Climatic microrefugia under anthropogenic climate change: implications for species redistribution. – Ecography 40: 253–266.

R Core Team 2017. R: A language and environment for statistical computing. – R Foundation for Statistical Computing, <https://www.R-project.org/>.

Riofrío-Dillon, G. 2013. Evolution of the acidity and nitrogen availability in the French forest soils over the 20th century – A spatiotemporal and multiscale approach based on the bioindicator character of plants. – Ph.D. thesis, AgroParisTech.

Riofrío-Dillon, G. et al. 2012. Toward a recovery time: forest herbs insight related to anthropogenic acidification. – Glob. Change Biol. 18: 3383–3394.

Robert, N. et al. 2010. France – In: Tomppo, E. et al. (eds), National Forest Inventories: pathways for common reporting. Springer Netherlands, pp. 207–221.

Scherrer, D. and Körner, C. 2011. Topographically controlled thermal-habitat differentiation buffers alpine plant diversity against climate warming. – J. Biogeogr. 38: 406–416.

ter Braak, C. F. and van Dame, H. 1989. Inferring pH from diatoms: a comparison of old and new calibration methods. – Hydrobiologia 178: 209–223.

Wagner, S. et al. 2011. Canopy effects on vegetation caused by harvesting and regeneration treatments. – Eur. J. For. Res. 130: 17–40.

## References

Belsley, D. A. et al. 2004. Regression Diagnostics: Identifying Influential Data and Sources of Collinearity. – John Wiley & Sons.

Dormann, C. F. et al. 2013. Collinearity: a review of methods to deal with it and a simulation study evaluating their performance. – Ecography 36: 27–46.

Fotheringham, A. S. and Oshan, T. M. 2016. Geographically weighted regression and multicollinearity: dispelling the myth. – J. Geogr. Syst. 18: 303–329.

Wheeler, D. and Tiefelsdorf, M. 2005. Multicollinearity and correlation among local regression coefficients in geographically weighted regression. – J. Geogr. Syst. 7: 161–187.

